# Plants emit informative airborne sounds under stress

**DOI:** 10.1101/507590

**Authors:** I. Khait, O. Lewin-Epstein, R. Sharon, K. Saban, R. Perelman, A. Boonman, Y. Yovel, L. Hadany

**Author notes:** Equal contribution (Y. Yovel, L. Hadany). Equal contribution (I. Khait, O. Lewin-Epstein, R. Sharon).

## Abstract

Stressed plants show altered phenotypes, including changes in color, smell, and shape. Yet, the possibility that plants emit *airborne sounds* when stressed – similarly to many animals – has not been investigated. Here we show, to our knowledge for the first time, that stressed plants emit airborne sounds that can be recorded remotely, both in acoustic chambers and in greenhouses. We recorded ∼65 dBSPL ultrasonic sounds 10 cm from tomato and tobacco plants, implying that these sounds could be detected by some organisms from up to several meters away. We developed machine learning models that were capable of distinguishing between plant sounds and general noises, and identifying the condition of the plants – dry, cut, or intact – based solely on the emitted sounds. Our results suggest that animals, humans, and possibly even other plants, could use sounds emitted by a plant to gain information about the plant’s condition. More investigation on plant bioacoustics in general and on sound emission in plants in particular may open new avenues for understanding plants and their interactions with the environment, and it may also have a significant impact on agriculture.

## Introduction

Plants exhibit significant changes in their phenotypes in response to stress. They differ visually, with respect to both color and shape, from unstressed plants [1-4]. They also emit volatile organic compounds (VOCs), e.g. when exposed to drought or herbivores [5, 6]. VOCs can also affect neighboring plants, resulting in increased resistance in these plants [7, 8]. Altogether, plants have been demonstrated to produce visual, chemical and tactile cues, which other organisms can sometimes respond to [9-12]. Nevertheless, the ability of plants to emit airborne sounds – that could potentially be heard by other organisms – has not been sufficiently explored [11, 13, 14].

Plants exposed to drought stress have been shown to experience cavitation – a process where air bubbles form, expand and explode in the xylem, causing vibrations [15, 16]. Yet, these vibrations have always been recorded by connecting the recording device directly to the plant [16, 17]. Such contact-based recording does not reveal the extent to which these vibrations could be sensed at a distance from the plant, if at all [17-19]. Thus, the question of airborne sound emission by plants remains unanswered [17, 20, 21].

Many animals, including herbivores and their predators, respond to sound [22-24]. Recently, plants were also demonstrated to respond to sounds [13, 25-27], e.g., by changing gene expression of specific genes [26, 27], or by increasing sugar concentration in their nectar [28]. If plants are capable of emitting informative airborne sounds, these sounds can potentially trigger a rapid effect on nearby organisms, including both animals and plants. Even if the emission of the sounds is entirely involuntarily, and is merely a result of the plant’s physiological condition, nearby organisms that are capable of hearing the sounds could use them for their own benefit.

We therefore set to examine whether plants emit informative airborne sounds, which may serve as potential signals or cues to their environment. Here we show for the first time that plants indeed emit airborne sounds, which can be detected from several meters away. Moreover, we show that the emitted sounds carry information about the physiological state of the plant. By training machine learning models we were able to achieve high accuracy in distinguishing between drought-stressed, cut, and control plants, based only on the sounds they emit. These results demonstrate the potential in studying phyto-acoustics and suggest that this form of communication may play an important role in plant ecology and evolution, and may have direct implications for plant monitoring in agriculture.

## Results

To investigate plants’ airborne sound emissions, we first constructed a reliable recording system, where the plants were recorded within an acoustic box, and then tested the system in a greenhouse.

### Plants emit airborne sounds when stressed

To investigate plants’ ability to emit airborne sound emissions, we first constructed a reliable recording system, in which each plant was recorded simultaneously with two microphones (see Fig. 1a for illustration, and Methods for details), within an acoustically isolated anechoic box. We recorded tomato (*Solanum lycopersicum*) and tobacco (*Nicotiana tabacum*) plants under different treatments – drought stress, cutting (of the stem), and controls. We focused on the ultrasonic sound range (20-150 kHz), where the background noise is weaker.

**Figure 1.**
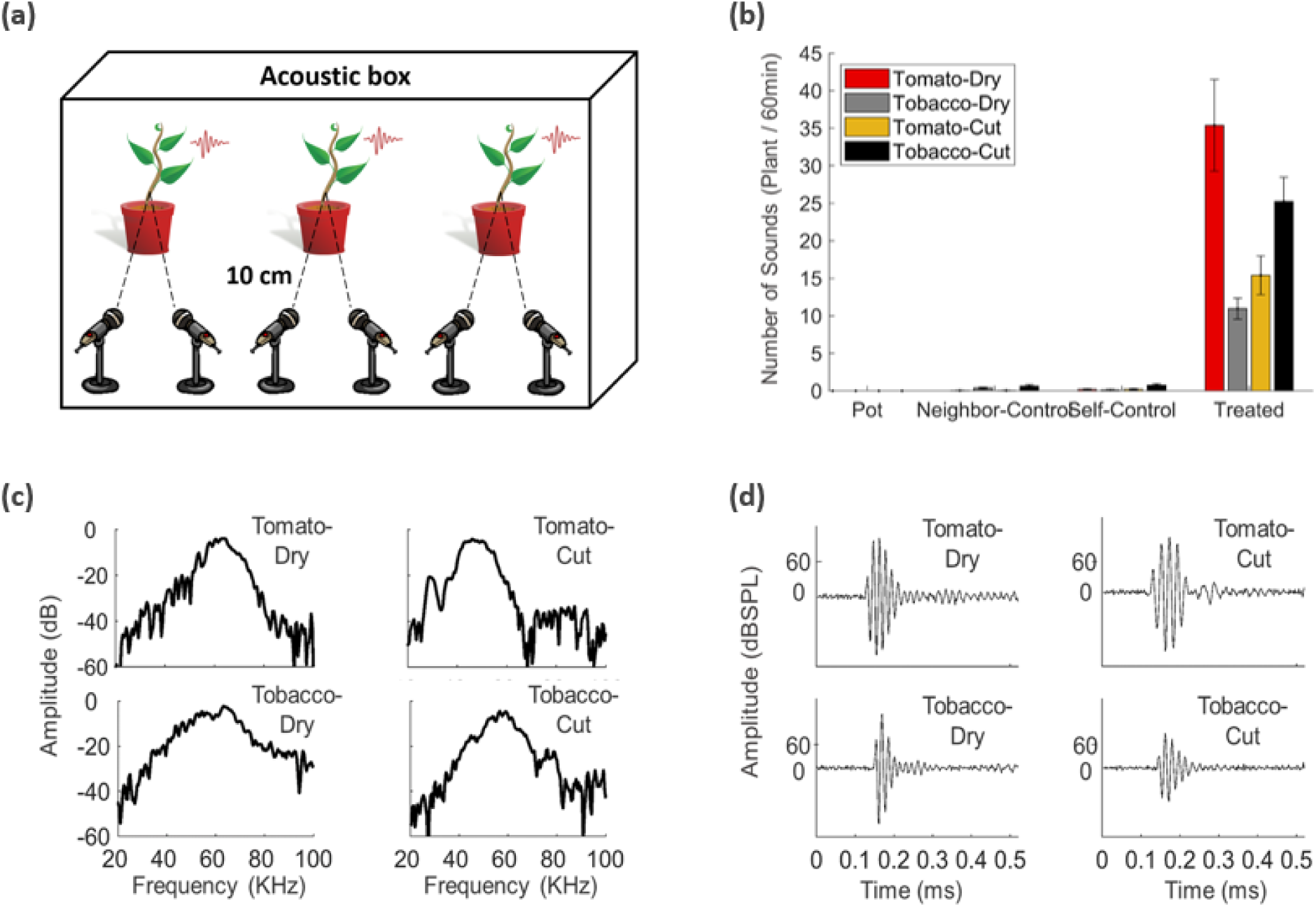
Plants emit remotely-detectable ultrasounds under stress. (a) Acoustic box setup. In each recording, three plants are placed inside an acoustic box with two directional microphones oriented at each plant. Using two microphones helps eliminating false detections resulting from electrical noise clicks of the recording system and cross-plant interference. (b) Mean number of sounds emitted during 60 minutes of recording by tomato and tobacco plants under two treatments, drought stress and cutting. Three control groups were used – empty pots, and two groups of untreated plants: self-control – the same plant before treatment, and neighbor-control – untreated plants that shared the acoustic box with treated plants. All treatment groups emitted significantly more sounds (p<e-6, Wilcoxon test with Bonferroni correction) than all control groups: self-control (Mean_self−control_ < 1 for all plant-treatment combinations) and neighbor control (Mean_neighbor−control_ < 1 for all plant-treatment combinations). The system did not record any sound from pots without plants during the experiments (Mean_pots_ = 0). 20 ≤ *n* ≤ 30 plants for each group. (c) Examples of time signals of sounds emitted by: a drought-stressed tomato, a drought-stressed tobacco, a cut tomato, and a cut tobacco. (d) The spectra of the sounds from (c).

We found that plants emit sounds, and that both drought-stressed plants (see Methods) and cut plants emit significantly more sounds than plants of any of the control groups (p<e-6, Wilcoxon test for each of the 12 comparisons with Holm-Bonferroni correction). Three controls were used for each plant species and treatment: recording from the same plant before treatment (*self-control*), recording from an untreated same-species neighbor plant (*neighbor-control*, see Methods), and recording an empty pot without a plant (*Pot*). The mean number of sounds emitted by drought-stressed plants was 35.4±6.1 and 11.0±1.4 per hour for tomato and tobacco, respectively, and Cut tomato and tobacco plants emitted 25.2±3.2 and 15.2±2.6 sounds per hour, respectively (Fig. 1b). In contrast, the mean number of sounds emitted by plants from all the control groups was lower than 1 per hour. Our system did not record any sound in the *Pot* control (Fig. 1b).

How does a stressed plant sound? Figs. 1c-d show examples of raw recorded time signals and their spectra as recorded from drought-stressed and cut plants. The mean peak sound intensity recorded from drought-stressed tomato plants was 61.6±0.1 dBSPL at 10 cm, with a mean peak frequency of 49.6±0.4 kHz (frequency with maximal energy), and the mean intensity recorded from drought-stressed tobacco sounds was 65.6±0.4 dBSPL at 10 cm, with a mean frequency of 54.8±1.1 kHz. The mean peak intensity of the sounds emitted by cut tomato plants was 65.6±0.2 dBSPL at 10 cm with a mean peak frequency of 57.3±0.7 kHz (frequency with maximal energy), and the mean intensity of the sounds emitted by cut tobacco plants was 63.3±0.2 dBSPL at 10.0 cm distance with a mean frequency of 57.8±0.7 kHz. The distributions of sound peak intensity and the maximum energy frequency of cut and drought-stressed plants are shown at Fig. 1c. Spectrograms of raw recorded sounds from cut and drought-stressed plants are shown at Supporting Information Fig. S1.

### The sounds of plants reveal their condition

To test whether we could identify different plant conditions and species based on their sound emissions, we trained a regularized machine learning classifier. We divided the sounds to four groups, with all the combinations of two plant types – tomato and tobacco, and two treatments – drought or cutting. The treatments were applied to the plants before the beginning of the recording. A binary classifier was trained to distinguish between two equal-size groups (“pair”) in each comparison (Tomato-Dry vs Tomato-Cut; Tobacco-Dry vs Tobacco-Cut; Tomato-Dry vs Tobacco-Dry; Tomato-Cut vs Tobacco-Cut). For cross validation, the model was tested only on plants that were not a part of the training process (see Methods for more details). We used a Support Vector Machine (SVM) as the classifier with several methods of feature extraction – basic [29, 30], MFCC [31], and a scattering network [23].

The SVM classifier with scattering network for feature extraction achieved the best results: ∼70% accuracy for each of the four pairs (Fig. 2b red line), significantly better than random (p<e-12 for each pair, Wilcoxon rank sum test with Holm-Bonferroni correction). The same classifier was trained to discriminate between the electrical noise of the system (see Methods) and the sounds emitted by the plants (Tomato vs Noise, Tobacco vs Noise) and achieved above 98% accuracy for both (Fig. 2b). The results were also robust to the dimension of the descriptors and the scattering network specific parameters (Fig. S2). Using the SVM classifier with MFCC as the input features, the results were still significantly better than random (p<e-4 for each pair, corrected as above), and even when we only used 4 basic features the results were significantly better than random for 5 of the 6 pairs (p<e-6 for each of them, adjusted; Fig. 2b). However, Scattering network performed better than either MFCC (p<0.05) or Basic (p<e-6) for all the pairs, using Wilcoxon sign rank test.

**Figure 2.**
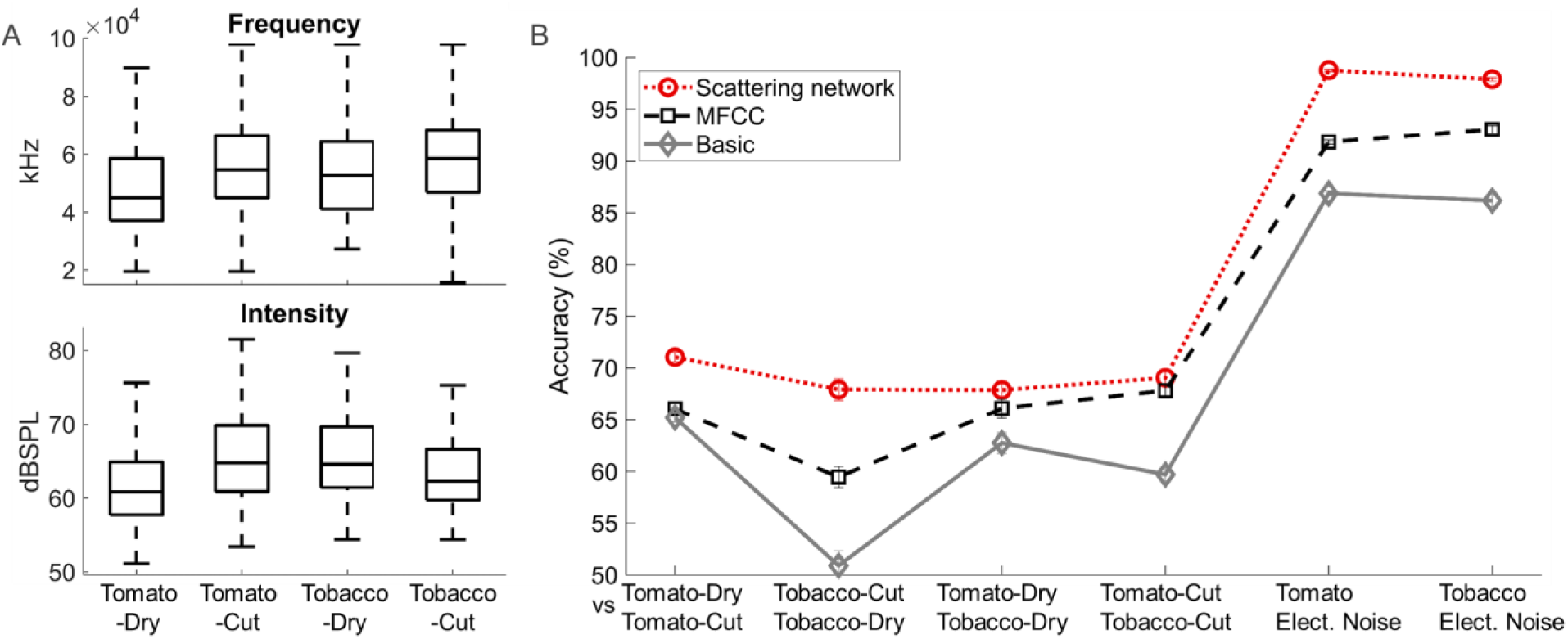
Plant condition and species can be detected from a distance by listening to its sound emissions. (a) The recorded sounds intensity peak and the max energy frequency for the four groups – drought stressed tomato plants, cut tomato plants, drought stressed tobacco plants and cut tobacco plants. (b) The accuracy of sound classification achieved by different feature extraction methods, with an SVM classifier. The best results were obtained using the scattering network method for feature extraction (red line, p < e-12 for each pair). Using MFCC for feature extraction the results were also highly significant (black dashed line, p<e-4 for each pair) and even basic methods for feature extraction allowed for better-than-random classification (gray line, p<e-6 for each pair apart from one case: Tobacco dry vs. Tobacco cut, which was bot significant with the basic method). The comparisons Tomato vs Elect. Noise and Tobacco vs Elect. Noise related to electrical noise of the system. Training set size of the two groups in each pair was equal (400 < sounds for each pair, see Table S2), and significance levels for each pair were calculated using Wilcoxon rank sum test with Holm–Bonferroni correction for multiple comparisons.

### Plant stress can be identified from its sounds in a greenhouse

Finally, we tested the acoustic behavior of plants in a more natural setting, a greenhouse. A major challenge we had to address was distinguishing between sounds generated by plants and the general noises in a greenhouse that were absent in the acoustic box (e.g. wind, rain, air-conditioning, construction). We first constructed a greenhouse noises library, by recording in the empty greenhouse, without any plant inside. We then trained a convolution neural network (CNN) model to distinguish between these greenhouse noises and the sounds of dry tomatoes recorded in the acoustic box (see details in Methods section). The model obtained from this training achieved 99.7% balanced accuracy in a cross-validation examination (see Figure 3 and figure S3).

**Figure 3.**
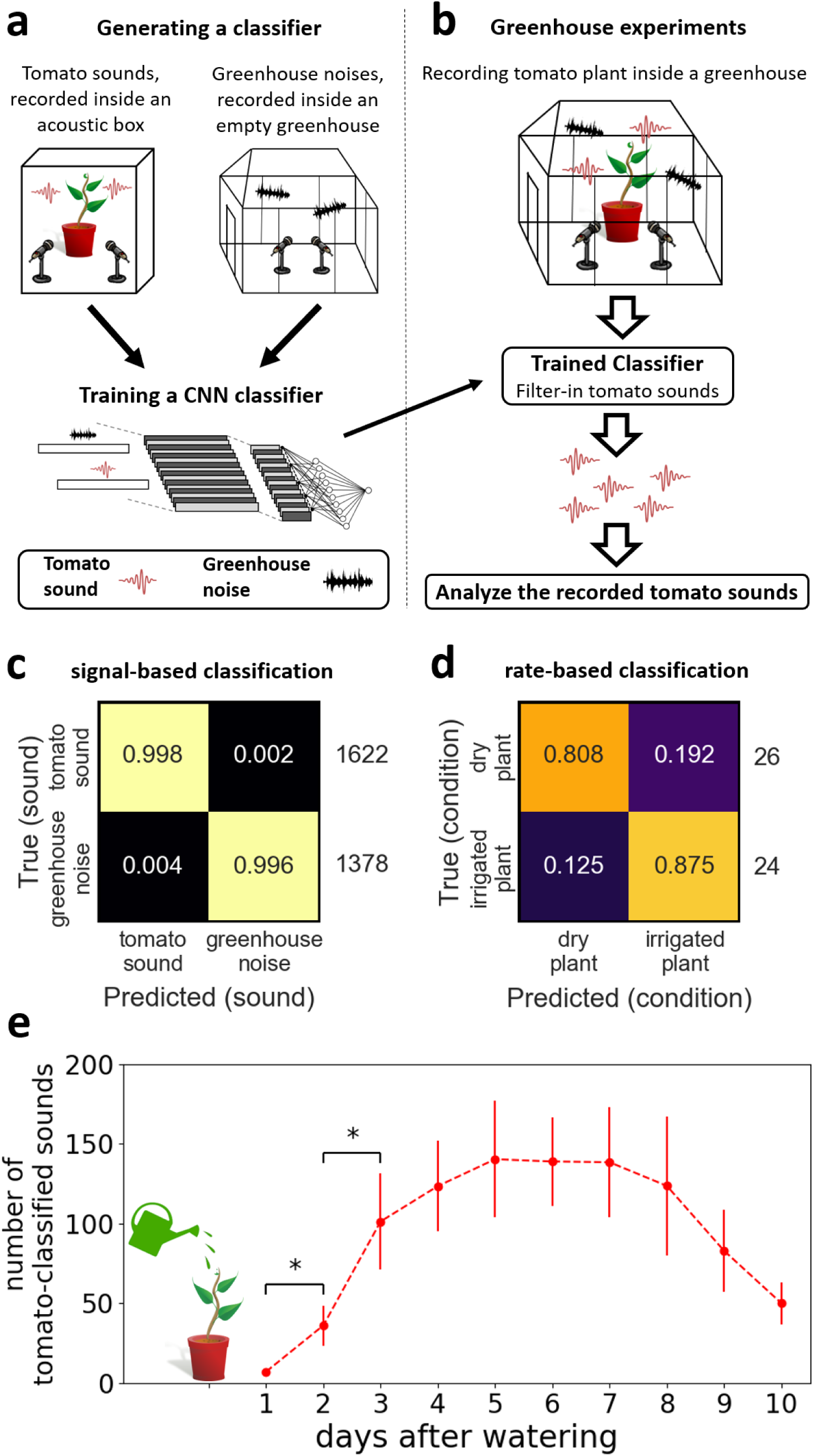
Acoustic detection of plant condition in the greenhouse. (a) Illustration of the procedure used to train a classifier that distinguishes between tomato sounds and greenhouse noises. A greenhouse noises library was first generated, by recording inside an empty greenhouse for several days. Using this library and the library of tomato sounds recorded in an acoustic box, we trained a convolution neural network (CNN) classifier to distinguish between tomato sounds and greenhouse noises. **(b) Illustration of the recordings in the greenhouse**. Tomato plants were recorded in the greenhouse. The recorded sounds were filtered using the trained CNN classifier, leaving only the tomato-classified sounds. **(c)** Confusion matrix showing the success of the trained CNN classifier in distinguishing between tomato sounds and greenhouse noises. Balanced accuracy score of ∼99.7%. **(d)** Confusion matrix showing the success in distinguishing between dry tomato plants and irrigated tomato plants, based on one hour of recording inside a greenhouse. The condition of the plant (dry / irrigated) was here decided based on the number of recorded tomato-classified sounds: if above three the plant was classified as “dry”, and otherwise as “irrigated”. Balanced accuracy score of ∼84% (p<e-5; Fisher exact test). **(e) The number of tomato-classified sounds per day during dehydration.** Tomato plants (*N* = 21) were recorded in the greenhouse for ten consecutive days without watering (starting one day after watering). The recorded sounds were then filtered using the trained CNN classifier, leaving only the tomato-classified sounds. We find significant difference in the amount of sounds between the following consecutive days: 1-2 and 2-3 (Wilcoxon signed-ranks tests, corrected for 9 comparisons between pairs of consecutive days using Holm-Bonferroni method; The ‘*’ markings in the figure represent adjusted p-values < 0.05).

After we succeeded filtering greenhouse noises, we could use “clean” plant sounds to classify plant condition in the greenhouse. Results from the acoustic box recordings showed that drought-stressed plants emitted significantly more sounds than control plants (Fig. 1). We thus used the number of tomato sounds emitted within an hour of recording to distinguish between drought-stressed plants and control plants, in the greenhouse. Each plant was recorded either one day after watering (control) or five days after watering (drought-stressed). This classification method, which only counted plant sounds, achieved ∼84% accuracy, significantly distinguishing between drought-stressed and control tomato plants (p<e-5, Fisher exact test; see further details in figures S4 and S5).

We continued to study the acoustic manifestation of the dehydration process, by recording tomato plants for ten consecutive days. Each tomato plant was watered, and then placed in the greenhouse for ten days of recording without watering. We used the CNN model mentioned above (figure 3a) to separate tomato sounds from greenhouse noises. We then counted the tomato-classified sounds recorded on each day from each tomato. These results revealed a consistent acoustic pattern: the plants emit very few sounds when irrigated, the number of sounds per day increases in the following 4-6 days, and then decreases as the plant dries (see figure 3e).

## Discussion

Our results demonstrate for the first time that plants emit remotely-detectable and informative airborne sounds under stress (Fig. 1). The plant emissions that we report, in the ultrasonic range of ∼20-100 kHz, could be detected from a distance of 3-5m (see Methods), by many mammals and insects (when taking their hearing sensitivity into account, e.g., mice [32] and moths [24]). Moreover, we succeeded in differentiating between sounds emitted under two different stress conditions – dry and cut – with accuracy of ∼70% using supervised machine learning methods (Fig. 2). Finally, we were able to filter plant sounds effectively from greenhouse noises (accuracy of ∼99.7%, Fig. 3b, c) and distinguish drought-stressed and control plants in a greenhouse, based only on the sounds they emit (accuracy of ∼84%, Fig. 3d, e). These findings can alter the way we think about the Plant Kingdom, which has been considered to be almost silent until now [20].

Our work can be extended in several ways. First, our results can be generalized to other species of plants from different families. In a preliminary study, we successfully recorded sounds from additional plants from different taxa, e.g., *Mammillaria spinosissima* cacti and *Henbit deadnettle* (Fig. S6). We thus expect that many plants have the ability to emit sounds, but the exact characteristics of these sounds, and the similarity between groups, are yet to be identified. Third, future studies could explore the sounds emitted under different plant states, including other stress conditions such as disease, cold, herbivores attack, or extreme UV radiation, and other life stages, such as flowering and fruit bearing. Once a large library of plant sounds is constructed, it could be analyzed by modern tools to obtain additional insights.

A possible mechanism that could be generating the sounds we record is cavitation – the process whereby air bubbles form and explode in the xylem [15, 16]. Cavitation explosions have been shown to produce vibrations similar to the ones we recorded [15, 16], but it has never been tested whether these sounds are transmitted through air at intensities that can be sensed by other organisms. Regardless of the specific mechanism generating them, the sounds we record carry information, and can be heard by many organisms. If these sounds serve for communication a plant could benefit from, natural selection could have favored traits that would increase their transmission.

We have shown that plant sounds can be effectively classified by machine learning algorithms. We thus suggest that other organisms may have evolved to classify these sounds as well, and respond to them. For instance, many moths – some of them using tomato and tobacco as hosts for their larvae [33, 34] – can hear and react to ultrasound in the frequencies and intensities that we recorded [22-24]. These moths may potentially benefit from avoiding laying their eggs on a plant that had emitted stress sounds. We hypothesize that even some predators may use the information about the plant’s state to their benefit. For example, if plants emit sounds in response to a caterpillar attack, predators such as bats [35] could use these sounds to detect attacked plants and prey on the herbivores [36], thus assisting the plant. The same sounds may also be perceived by nearby plants. Plants were already shown to react to sounds [13, 25-28] and specifically to increase their drought tolerance in response to sounds [37, 38]. We speculate that plants could potentially hear their drought stressed or injured neighbors and react accordingly.

Finally, plant sound emissions could offer a novel way for monitoring crops water state – a question of crucial importance in agriculture [39]. More precise irrigation can save up to 50% of the water expenditure and increase the yield, with dramatic economic implications [39, 40]. In times when more and more areas are exposed to drought due to climate change [41], while human population and consumption keep increasing [42], efficient water use becomes even more critical, for both food security and ecology. Our results, demonstrating the ability to distinguish between drought-stressed and control plants based on plant sounds, open a new direction in the field of precision agriculture.

We demonstrated for the first time that stressed plants emit remotely detectable sounds, similarly to many animals, using ultrasound clicks not audible to human ears. We also found that the sounds contain information, and can reveal plant state. The results suggest a new modality of signaling for plants and imply that other organisms could have evolved to hear, classify and respond to these sounds. We suggest that more investigation in the plant bioacoustics field, and particularly in the ability of plants to emit and react to sounds under different conditions and environments, may reveal a new pathway of signaling, parallel to VOCs, between plants and their environment.

## Materials and Methods

### Plants materials and growth conditions

Tomato – *Solanum lycopersicum* ‘Hawaii 7981’ [43] – and tobacco – *Nicotiana tabacum* ‘Samsun NN’ – were used in all the experiments. All the plants were grown in a growth room at 25 °C and kept in long-day conditions (16 h day, 8 h night). The plants were tested in the experiments 5-7 weeks after germination.

### Recording protocol

#### In the acoustic box

The recordings were performed in a 50 × 100 × 150 *cm*^3^ acoustically isolated box tiled with acoustic foam on all sides to minimize echoes. Two cable holes, 2 cm radius each, were located in two corners of the box and covered with PVC and acoustic foam. Inside the acoustic box were only the recorded plants, 6 microphones connected to an UltraSoundGate 1216H A/D converter (Avisoft). The PC and all the electricity connections were in the room outside the acoustic box. Two USB cables connected the PC to the 1216H device inside the box, through the holes. There was no light inside the acoustic box.

The recordings were performed using a condenser CM16 ultrasound microphone (Avisoft), digitized using an UltraSoundGate 1216H A/D converter (Avisoft), and stored onto a PC. The sampling rate was 500 KHz per channel. We filtered the recordings above 20 KHz. A recording started only when triggered with a sound which exceeded 2% of the maximum dynamic range of the microphone. Two microphones were directed at each plant stem, from a distance of 10 cm. Only sounds that were recorded by both microphones were considered as a “plant sound” in the analysis afterwards. The frequency responses of the microphones can be found in Avisoft website.

#### In the greenhouse

The recordings were performed in a greenhouse in Tel-Aviv University. Inside the greenhouse were only the recorded plants. The recordings were performed using the same hardware and setting, as mentioned in the acoustic box recording section, apart from the acoustic box itself.

### Data pre-processing

Data processing was performed off-line using matlab code we developed (MATLAB 8.3, The MathWork Inc.), with the following steps: 1. Identifying the microphone that had recorded the highest intensity peak at the moment the recording started. 2. Selecting the sounds that were detected by two microphones oriented at the same plant at the same time, and saving them for further analysis. The recordings produce 6 channel wav files. A processed recording includes a short section of 1ms before and 1ms after the peak of the recorded sound that triggered the system to record. “Greenhouse noise” sounds were obtained when the Greenhouse included only acoustic equipment without plants or pots, by the two microphones (thus not including “Electrical noise”).

### Acoustic-box Drought stress experiment

Each plant was recorded twice: first before drought treatment (“self-control”), and again after it. In the first recording, all the plants were healthy and their soil was moist. Then, for 4-6 days, half of the plants were watered while the other half were not, until the soil moisture in the pots of un-watered plants decreased below 5%. Then, the plants were recorded again at the same order. In each recording session three plants were recorded simultaneously for one hour and each triplet of plants included at least one watered and one un-watered plant to allow “neighbors-control” – watered plants that were recorded while sharing the acoustic box with un-watered plants. Soil moisture content was recorded using a hand-held digital soil moisture meter - Lutron PMS-714.

### Acoustic-box Cut stress experiment

The experiment followed the experimental design of the drought stress experiment described above, but drought stress was replaced with cutting of the plant. Here the pot soil was kept moist for all the plants throughout the experiment. The plants included in the treatment group were cut with scissors close to the ground right before the recording started. The severed part of the plant, disconnected from the roots, was recorded. We used the same controls of the drought stress experiment.

### Greenhouse experiment

In each recording session three plants were recorded simultaneously for one hour. All the recorded plants were grown in a growth room, and were brought to the greenhouse only for the recording session. Each plant was recorded either one day after irrigation (control plants) or 4-6 days after irrigation (drought-stressed plants).

### Classifying sounds recorded in the acoustic box

Our classification method was composed of two main stages. First, we extracted various acoustic features from the raw recorded signals (see Data Pre-processing section). Second, we trained a model to classify plant sounds into classes based on the feature representation obtained in the first stage. We used three methods of feature extraction: (a) Deep scattering Network, as described in Andén and Mallat [44], red dotted line in Fig. 2b. This method extends MFCC while minimizing information loss. We used the implementation by ScatNet [45], with Morlet wavelets. The results were robust to the dimension of descriptors and the scattering network specific parameters: number of layers used; time support of low pass filter; and Q-Factor (Fig. S2). The values of the specific parameters used in this work are shown at Table S1. (b) MFCC feature extraction (dashed black line in Fig. 3b). We used the Ellis Dan implementation [31]. (c) Basic features. The basic features we used were energy, energy entropy, spectral entropy, and maximum frequency (gray line in Fig. 2b) [29, 30]. We used SVM with Radial kernel with the LIBSVM implementation as classifier. After feature extraction, we used Z-score for normalization of the features and PCA to reduce the dimensionality of the problem. We used only the training set to choose the number of features.

During the training process, we leave all the emitted sounds of one plant out for cross validation. Then we constructed the training set such that the two compared groups would be of the same size. We repeated the process so that each plant constructed the testing group exactly one time. The accuracy of the classification was defined as the percentage of correct labeling over the total size of the testing set [46, 47]. The numbers of plants in each group are shown at the Table S3.

### Classifying sounds recorded in the greenhouse

For classification of greenhouse noise and drought-stressed tomato sounds, we used a convolution neural network implemented using Python 3.6 with keras package and tensorflow backend. The network receives the vectors of the processed recordings (2ms long time signals, see Data pre-processing section) as input. The network is composed of three 1-D convolution layers, each followed by a maxpooling layer, and one fully-connected layer afterwards. At the end, we add a layer of size 1 with sigmoid activation. The model is trained with binary crossentropy loss function and ‘adam’ optimizer. This model achieved a balanced accuracy score of ∼0.997 over leave-one-plant-out cross validation (namely in each iteration we trained a model on the sounds from all plants except one plant, and then we validated the model on this latter plant).

### Statistical analysis

For statistical analysis of the number of sound emissions for the treatment and the control groups (Fig. 1b) we used the Wilcoxon rank-sum test. To compare our classifier to random result (Fig. 2b), we used the binomial probability distribution function (PDF) and calculate the probability to get the classifier accuracy or higher randomly for each group. To compare the results obtained when using scattering network for feature extraction to the results obtained when using MFCC or basic feature extraction methods (Fig. 2b), we used Wilcoxon sign rank test with Holm-Bonferroni correction. To test the success in distinguishing between drought-stressed and control plants (Fig. 3) we used Fisher’s exact test.

## Acknowledgements

We thank Daniel Chamovitz, Gal Chechik, Tuvik Beker, and Judith Berman for comments on the paper; Guido Sessa, Doron Teper, Guy Sobol, Yura Pupov, Rotem Shteinshleifer, Odelia Pisanty, Eilon Shani, and Meirav Leibman-Markus for helping with plants materials; Yoel Shkolnisky, Marine Veits, Ilia Raysin, Uri Obolski, Yoav Ram, Eyal Zinger, Yael Gurevich, Eylon Tamir, Yuval Sapir, Yaara Blogovski and Ruth Cohen-Khait for comments on the way. The Titan Xp used for this research was donated by the NVIDIA Corporation.

## Author Contributions

LH and IK conceived the study. LH, YY and IK, designed the research. IK, RP, KS and OLE performed the experiments. RS, OLE, IK, AB and LH analyzed the data. YY and LH supervised the experiments. IK, OLE and RS contributed equally to the study. LH and YY contributed equally to the study. All authors discussed the results and took part in writing the manuscript.

## Data and materials availability

The data will be deposited on Dryad upon acceptance.

## Supporting Information

**Fig. S1** Examples for spectrograms of sounds which emitted by stressed plants.

**Fig. S2** Comparison of different scattering network configurations.

**Fig. S3** ROC curves of four neural network models.

**Fig. S4** Tomato sounds distribution.

**Fig. S5** ROC curve of plant condition decision threshold.

**Fig. S6** Recorded sounds from different plants.

**Table S1** Parameters used in the feature extraction phase.

**Table S2** Pairs total sizes.

**Table S3** Groups sizes.

