## Supplementary information for "Plants emit informative airborne sounds under stress"

**This PDF file includes:**

Figures S1 to S6

Tables S1 to S3

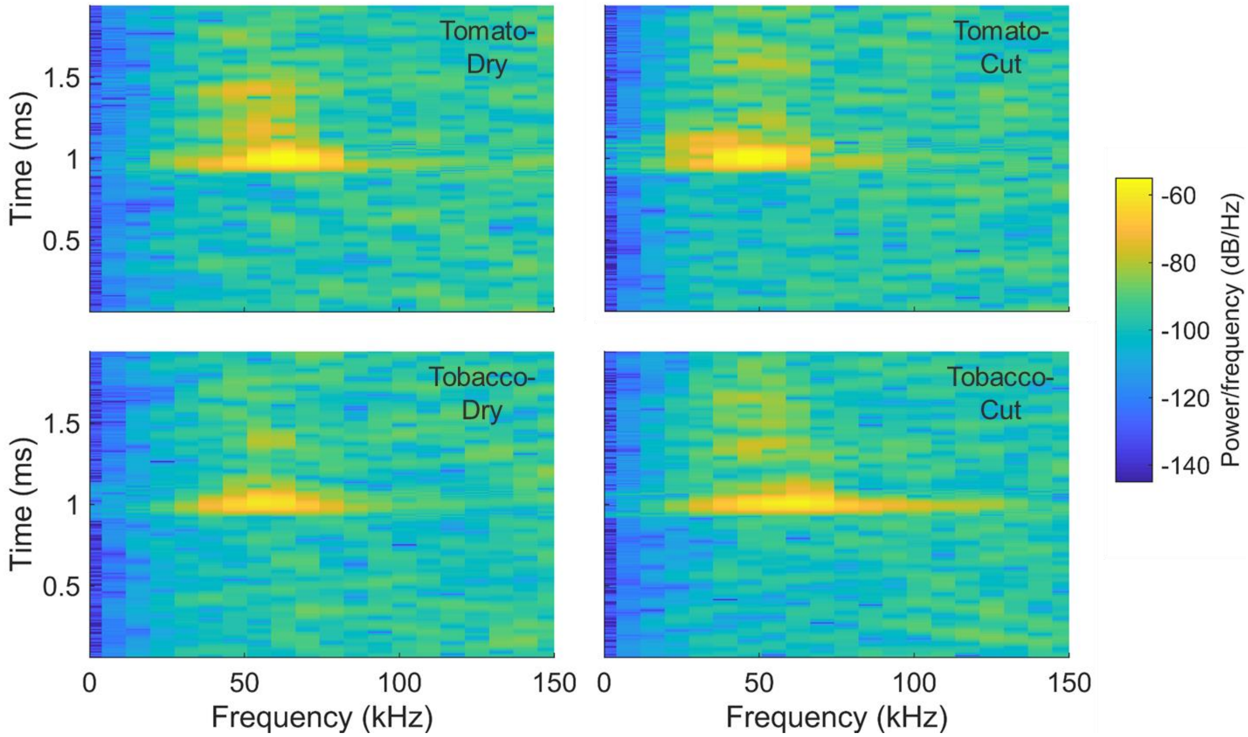


**Figure S1. Examples for spectrograms of sounds which emitted by stressed plants.** Examples for spectrograms of sounds which time signal and spectra are showed at Fig. 2**b, c.** The sounds were emitted by: a drought stressed tomato, a drought stressed tobacco, a cut tomato and a cut tobacco.

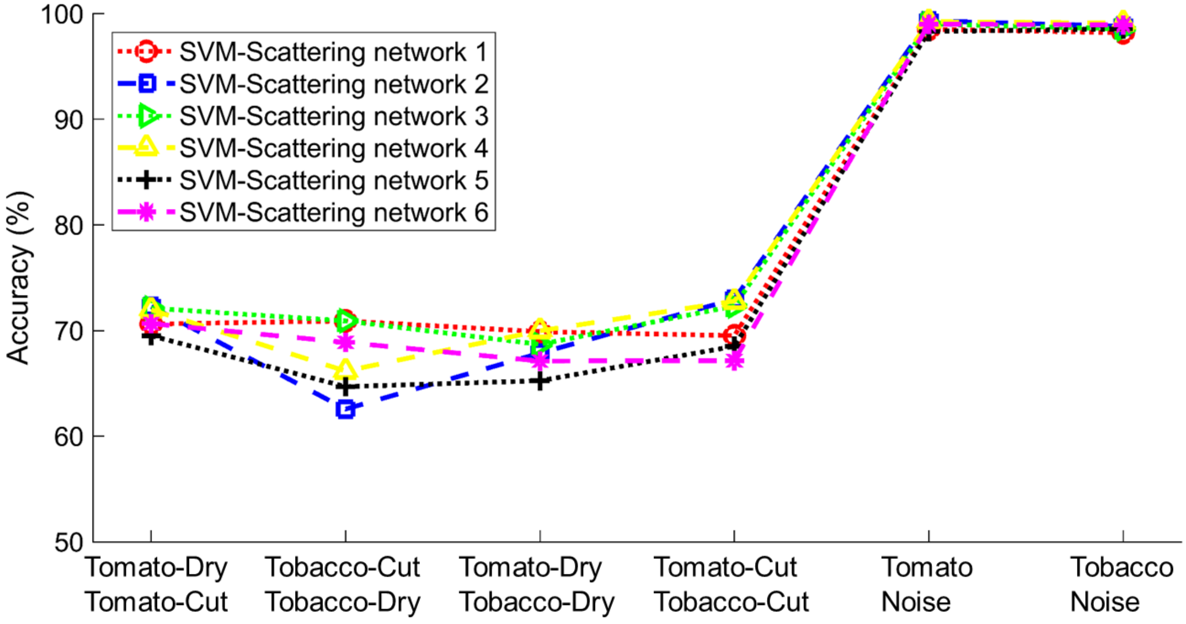


**Figure S2. Comparison of different scattering network configurations.** The accuracy of sound classification with 6 different configurations for the scattering network, using SVM as classifier. Each line represents a different feature set, all obtained by scattering network. The scattering networks had different time intervals, different Q-factors, and a different number of filters, for exact values see Table S1.

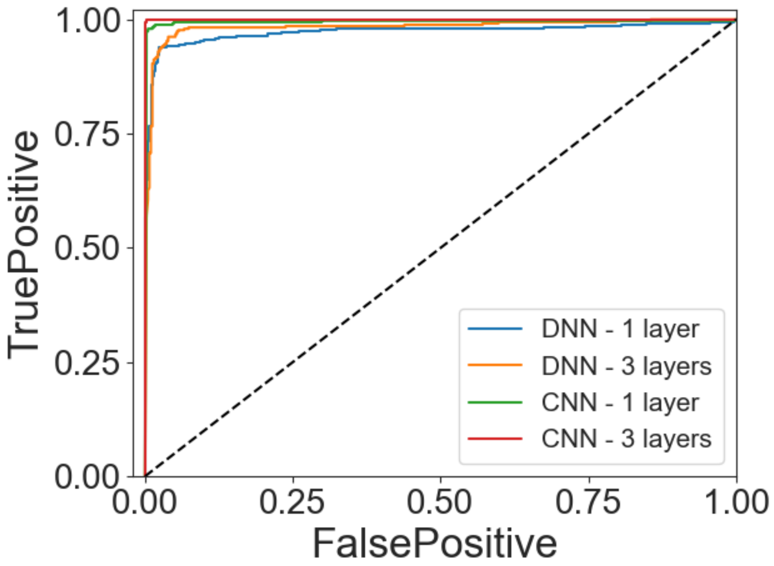


**Figure S3. Receiver operating characteristic (ROC) curves presenting the performance of four different neural network models, with respect to distinguishing between tomato sounds and greenhouse noises.** We plot the true positive rate as function of the false positive rate of four different neural network models: a model with one fully-connected layer (size 128), a model with three fully-connected layers (sizes 32, 64 and 128), a convolution network with one layer (depth 128), and a convolution network with three layers (depths 32, 64, 128). We do so for a model for classification of tomato sounds and greenhouse noises. The area under the curve score (AUC) for all models (same order as mentioned above): 0.97287, 0.98325, 0.99452, 0.99999. These results were obtained by training the model on 75% of the data and testing on the rest 25%.

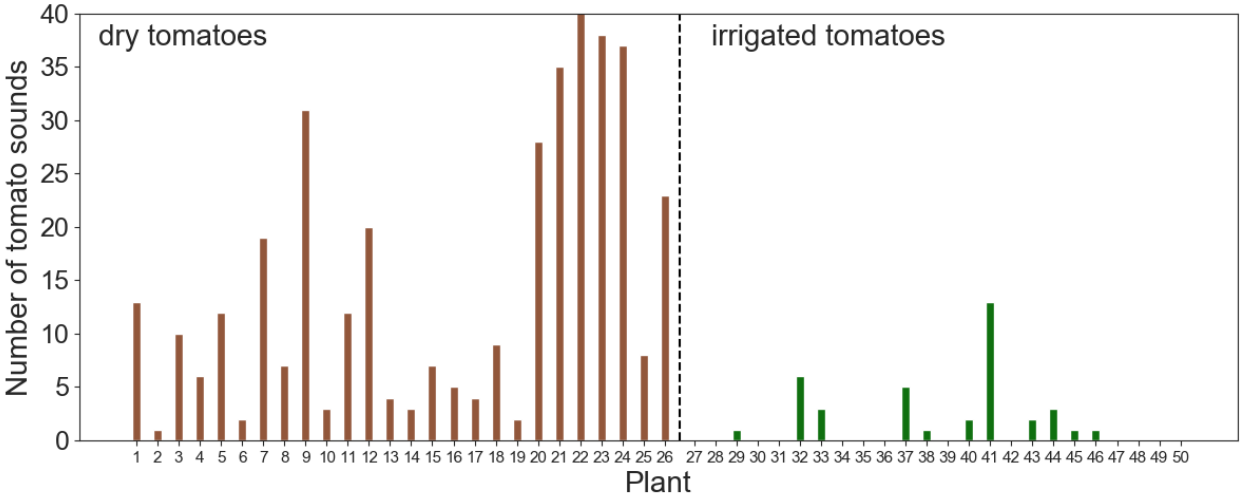


**Figure S4. Tomato-classified sounds per hour.** We recorded dry and irrigated tomato plants for one hour in the green house. Then, using a trained Convolutional Neural Network (CNN) we filtered the greenhouse noises and left only the tomato classified sounds. We here show the number of tomato-classified sounds, recorded during one hour in the greenhouse for dry and irrigated tomato plants. The y-axis is truncated at 40 for better resolution; plant 22 emitted 95 sounds.

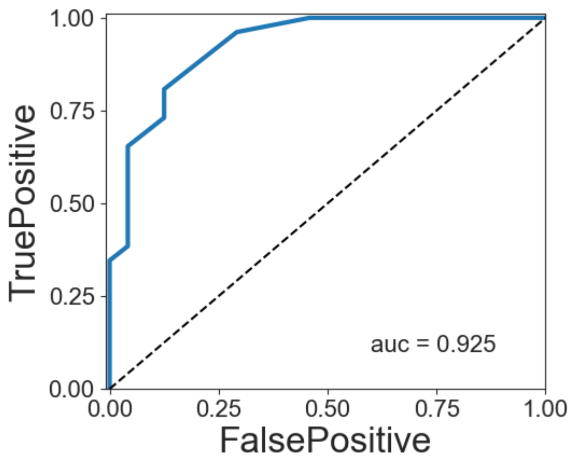


**Figure S5. Receiver operating characteristic (ROC) curves presenting the performance of the classification model used to distinguish between dry and irrigated tomato plants.** We plot the true positive rate as function of the false positive rate for all possible decision threshold, where a decision threshold is of the type: above a certain amount $(n_c)$ of tomato-classified sounds per hour we classify the plant as dry, otherwise we classify the plant as irrigated.

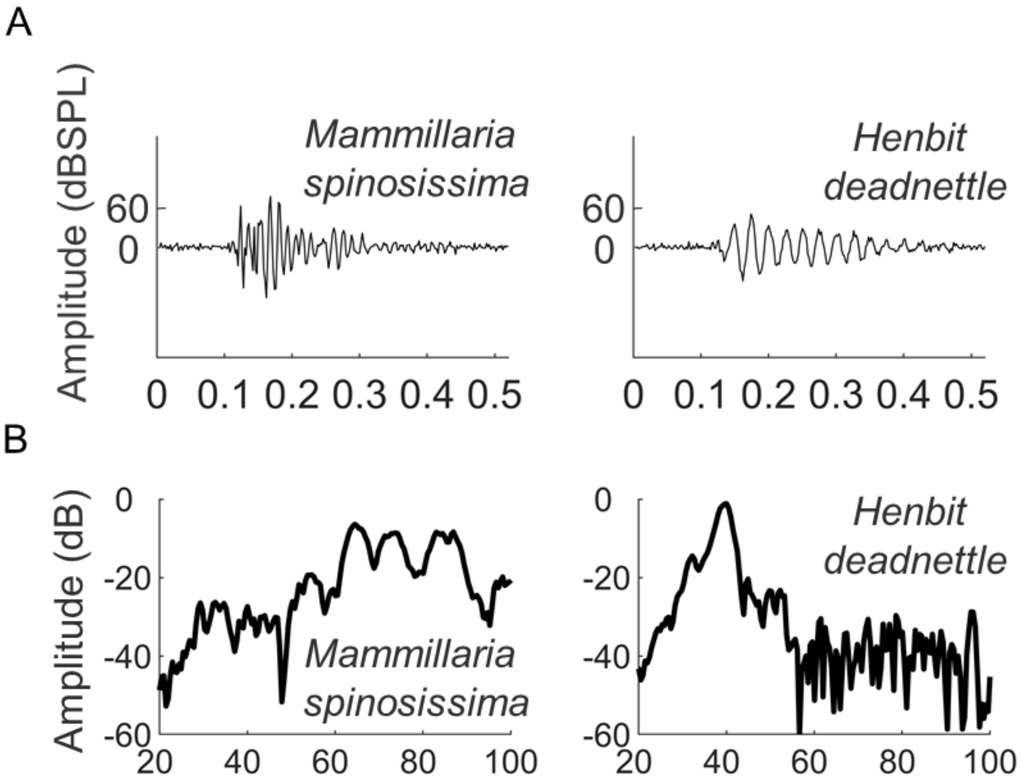


**Figure S6. Recorded sounds from different plants** (a) Examples of time signals of sounds emitted by *Mammillaria spinosissima* cacti and *Henbit deadnettle* plant. (b) The spectra of the sounds from (a).

**Table S1- Parameters used in the feature extraction phase**

| Method | Normalization | PCA | Parameters |
|---|---|---|---|
| Basic | - | - | - |
| MFCC | - | - | - |
| SN 1 | Z-Score | 120 | M = 2, T = 1ms, Q = [12] |
| SN 2 | Z-Score | - | M = 2, T = 2ms, Q = [1 1] |
| SN 3 | Z-Score | - | M = 1, T = 3ms, Q = [8] |
| SN 4 | - | 80 | M = 1, T = 3ms, Q = [14] |
| SN 5 | Z-Score | - | M = 2, T = 1ms, Q = [12 2] |
| SN 6 | Z-Score | - | M = 1, T = 1ms, Q = [1] |

Scattering Network is denoted SN. PCA column describes the number of principle component used. The parameters M, T, and Q refer to Scattering Network parameters. M = number of layers used, T = time support of low pass filter, and Q = Q-Factor. When written in the text SN without any number we use the configuration of SN1.

**Table S2- Pairs total sizes (2 equal-size groups in each pair)**

| Pair | | |
|---|---|---|
| Group 1 | Group 2 | Total number of sounds in pair |
| Tomato - Dry | Tomato - Cut | 1262 |
| Tobacco - Dry | Tobacco - Cut | 418 |
| Tomato - Dry | Tobacco - Dry | 418 |
| Tomato - Cut | Tobacco - Cut | 932 |
| Tomato - All | Noise | 3868 |
| Tobacco - All | Noise | 1350 |

**Table S3- Groups sizes**

| Group | Recorded sounds (#) | Individual plants (#) | Median number of sounds emitted by one plant (#) |
|---|---|---|---|
| Tomato-Dry | 1612 | 51 | 21 |
| Tomato-Cut | 631 | 40 | 11.5 |
| Tobacco-Dry | 209 | 23 | 8 |
| Tobacco-Cut | 466 | 19 | 24 |
| Noise | 1934 | - | - |
